# Naturally Occurring Shorter and Tumor-associated Mutant RNF167 Variants Facilitates Lysosomal Exocytosis and Plasma Membrane Resealing

**DOI:** 10.1101/772731

**Authors:** Sreeja V. Nair, Nikhil Dev Narendradev, Rithwik P. Nambiar, Rakesh Kumar, Srinivasa M. Srinivasula

## Abstract

Lysosomal exocytosis and resealing of damaged plasma membrane play critical roles in physiological and pathological processes, including, restoration of cellular homeostasis and tumor invasion. However, to-date, only a few regulatory molecules of these biological processes have been identified. Moreover, no mutations in any of the known regulators of lysosomal exocytosis in primary tumors of patients have been characterized. Here we demonstrate that RNF167, a lysosomal associated ubiquitin ligase, negatively regulates lysosomal exocytosis by inducing perinuclear clustering of lysosomes. Importantly, we also characterized a set of novel natural mutations in RNF167, which are commonly found in diverse tumor types. We found that RNF167-K97N mutant, unlike the wild-type, localizes in the cytoplasm and does not promote perinuclear lysosomal clustering and that cells expressing RNF167-K97N exhibit dispersed lysosomes, increased exocytosis, and enhanced plasma membrane repair. Interestingly, these functional features of RNF167-K97N were shared with a naturally occurring short version of RNF167, i.e. isoform b. In brief, the results presented here reveal a novel role of RNF167 as well as its natural variants, RNF167-K97N and RNF167-b as an upstream regulator of lysosomal exocytosis and plasma membrane resealing which might play an important role in organelle dynamics or tumor progression or both.

## Introduction

Lysosomes are specialized organelles with an inherent function in recycling and degrading the cargo delivered to them through endocytosis, phagocytosis, and autophagy. In recent years, lysosomes have emerged as dynamic structures regulating physiologic and pathologic cellular processes such as antigen presentation, destroying intracellular pathogens, plasma membrane repair, tumor invasion and metastasis, apoptotic cell death, metabolic signaling, and gene regulation (1). These diverse functions of lysosomal vesicles are primarily facilitated by their positioning in the cells and bidirectional movement between the plasma membrane and perinuclear region (2). The lysosomal movement towards the plus-end of microtubules (anterograde) and minus-end (retrograde) is mediated by kinesins and dynein motor proteins (3, 4) respectively. These motors generally do not bind lysosomes directly instead, the interaction is mediated by small GTPase like Rab7, Arl8b, and their effector proteins and lipids (2).

Under basal conditions, the majority of the lysosomes are concentrated in the center, known as perinuclear cloud and a few in the cell periphery (5). The distribution and motility of lysosomes are subjected to change depending on various conditions. For instance, cytosolic acidification promotes anterograde movement whereas alkalinization favors retrograde transport of lysosomes (6, 7). In general, perinuclear accumulation of lysosomes occurs during starvation (8), aggresome formation (9), drug-induced apoptosis (10), expression of pathogenic mutant forms of huntingtin (11) or leucine-rich repeat kinase 2 (LRRK2) (12), and lysosomal storage diseases (13, 14). Several protein complexes have been shown to regulate the intracellular positioning and movement of lysosomes. For example, the anterograde movement of lysosomes is mediated by a multisubunit complex, BLOC-1 related complex (BORC) which recruits Arl8b and its effector molecule SKIP (15, 16). Whereas retrograde movement is mediated by Rab7 along with its effector molecule RILP (17). Lysosomal movement and positioning play a decisive role in maintaining cellular homeostasis, dysregulation of which is a common event in various pathological conditions such as cancer and neurodegenerative diseases (1).

Extracellular acidification, a tumor microenvironment stimulus, promotes redistribution of lysosomes towards the cell periphery and this anterograde movement of lysosomes facilitates cancer growth, invasion, and metastasis (1). During invasion through the dense extracellular matrix of neighboring tissue spaces or metastasis tumor cells are prone to plasma membrane damages (18). Tumor cells cope up with these kinds of stresses by upregulating plasma membrane resealing (PMR) by lysosomal exocytosis. The damaged plasma membrane triggers an elevation in intracellular Ca^2+^ level resulting in anterograde movement of lysosomes. This peripheral pool of lysosomes docked to the cell surface, fuse with the plasma membrane and contributes to the resealing of the damaged plasma membrane (19). It has been shown that S100A11, an annexin binding protein, overexpressed in various tumors, enhances plasma membrane resealing in metastatic cells thereby protects them from plasma membrane stresses (18). Importantly, inhibition of lysosomal exocytosis could reverse invasiveness and chemoresistance in aggressive sarcoma cells (20). Inhibitors of sodium-proton exchangers (NHEs) have been shown to be very effective in preventing cell-surface directed lysosome trafficking, thereby decreasing lysosomal exocytosis and cell invasion (21), underlining the importance of lysosomal exocytosis in tumor progression.

Recent studies on RNF26, an endoplasmic reticulum (ER) associated ubiquitin ligase demonstrated the role of ubiquitin ligases in late endosomal (LE) positioning (5). It has been reported that ER localization and RING activity of the protein is essential for the perinuclear clustering of LE compartments. Another membrane associated ubiquitin ligase is RNF167. Though some reports suggested of RNF167 association with early endosomes (22), we and others found that it mostly localizes to lysosomes (23), with its overexpression inducing perinuclear clustering of lysosomes upon acetate Ringer’s solution treatment (24). This ubiquitin ligase consists of N-terminal signal peptide (SP), a protease associated (PA) domain, a transmembrane (TM) domain followed by a really interesting new gene (RING) domain (23, 25, 26). It has been shown that the PA domain is essential for the endosomal localization (22), while the RING domain confers ubiquitin ligase activity of the protein (22, 23). Examination of the NCBI database suggested the existence of different isoforms of RNF167, with one of the isoforms (RNA167-b), lacking the first 35 amino acids containing a putative signal peptide. Thus far, no evidence for the expression of any of the isoforms or their characterization is available. In this study, we provide evidence to establish that RNF167-a, is a negative regulator of lysosomal exocytosis. In contrast, RNF167-b and a naturally occurring tumor-associated mutant enhance lysosomal exocytosis and plasma membrane resealing. In summary, we report here for the first time how tumor-associated mutations in a regulator of lysosomal exocytosis contribute to tumor progression.

## Materials and Methods

### Cell culture and Stable cell preparation

HeLa and HEK-293T cells were cultured in Dulbecco’s modified Eagle’s medium (DMEM with 4.5g/l glucose, L-glutamine, 3.7 g/l sodium bicarbonate and sodium pyruvate) (AL007A, HiMedia, Mumbai, Maharashtra, India) supplemented with 10% FBS (10270-106, GIBCO-Thermo Fisher Scientific, Waltham, MA, USA) and 1% penicillin-streptomycin (A001A, HiMedia, Mumbai, Maharashtra, India). All cells were kept at 37°C, 5% CO_2_ incubator and all are obtained from ATCC.

HeLa cells stably expressing RNF167 WT was generated as follows: HEK-293T cells were transfected with 1:1:2 ratio of pCAG-VSVG, pUMVC, and pMYs-IRES-Puro RNF167 WT plasmids using Lipofectamine 3000 reagent (L3000-015, Invitrogen, Carlsbad, CA, USA) as per the manufacturer’s instructions. 4 days post-transfection, retrovirus was harvested and used to infect HeLa cells in the presence of 4 μg/ml polybrene. Stable cell lines were selected using 1μg/ml puromycin selection. Other stable cell lines mentioned in this study were also prepared using a similar protocol. Positive colonies were passaged further and used for experiments.

### Recombinant DNA constructs

The RNF167Δ1-35 construct was generated by cloning the coding sequence of RNF167 obtained by PCR amplification with primers:

FP-5’-GAAACTCGAGATGGACTTTGCAGACCTTCCAGC-3’ and

RP-5’-AAAGGGCCCGGACCAGGATAACAGGGGAAG-3’, followed by in-frame cloning into XhoI-ApaI sites of pEYFP-N1 (Clontech, Palo Alto, CA, USA). RNF167 WT and all the variants were cloned into pMYs-IRES-Puro (Cell Biolabs, San Diego, CA, USA) using Infusion cloning kit (Takara Bio Inc., Otsu, Japan) with primers:

FP-5’-GGTGGTACGGGAATTATGCACCCTGCAGCCTTC-3’ and

RP-5’-ATTTACGTAGCGGCCTTAGACCAGGATAACAGGG-3’. For RNF167Δ1-35:

FP-5’-GGTGGTACGGGAATTATGGACTTTGCAGACCTTC-3’ with the same reverse primer as mentioned above. All primers used in this study were synthesized and bought from Sigma Aldrich (St. Louis, MO, USA). The following plasmids were obtained from Addgene: GFP-rab11 WT was a gift from Richard Pagano (Addgene # 12674) (27), mTFP1-Lysosomes-20 was a gift from Michael Davidson (Addgene # 55495) (28), GFP-EEA1 WT was a gift from Silvia Corvera (Addgene # 42307) (29), pCAG-VSVG was a gift from Arthur Nienhuis & Patrick Salmon (Addgene # 35616), pUMVC was a gift from Bob Weinberg (plasmid # 8449) (30). A mammalian expression plasmid encoding GFP-Dynamitin (31) was a gift from Dr. Roberto Botelho (Ryerson University, Canada). Plasmids encoding C-terminal GFP fusion of RNF167 C268R, RNF167 R277L, RNF167 L329P, and RNF167ΔPA (22) were generously provided by Dr. Ruth H Palmer (University of Gothenburg, Sweden).

Amino acid substitutions in RNF167 were made using site-directed mutagenesis by Prime Star GXL-DNA polymerase (R050A, Takara Bio Inc., Otsu, Japan) according to the manufacturer’s protocol with the following primers:

5’-GCAACTTTGACCTCAATGTCCTAAATGCCCAG-3’and

5’-CTGGGCATTTAGGACATTGAGGTCAAAGTTGC-3’for RNF167 K97N.

5’-GTTGTATCCAGCACCAGAAACGGCTCCAGCG-3’

5’-CGCTGGAGCCGTTTCTGGTGCTGGATACAAC-5’for RNF167 R200Q.

5’-GATGTCTGTGCCATTAGCCTGGATGAATATGAG-3’and

5’-CTCATATTCATCCAGGCTAATGGCACAGACATC-3’ for RNF167 C233S.

### Antibodies and Reagents

The following antibodies were used in this study: rabbit anti-LAMP1(ab24170, Abcam, Cambridge, MA, USA), mouse anti-GFP (632375, Takara Bio Inc., Otsu, Japan), mouse anti-FLAG (F3165, Sigma Aldrich, St. Louis, MO, USA), mouse anti-LAMP1 (H4A3, Developmental Studies Hybridoma Bank, Iowa City, IA, USA), Cy5-AffiniPure Donkey Anti-Mouse IgG (715-175-150, Jackson Immuno Research Laboratories, Inc., West Grove, PA, USA), Goat anti-Rabbit IgG (H+L) Alexa Fluor 568 (A-11036), Goat anti-Mouse IgG (H+L) Alexa Fluor 488 (A-11001), Goat anti-Mouse IgG (H+L) Alexa Fluor 568 (A-11031). All the Alexa fluorophore-conjugated antibodies were purchased from Thermo Fisher Scientific (Rockford, IL, USA).

Reagents: Ionomycin (I3909), DMSO (D8418), Streptolysin-O (SAE0089), Propidium iodide (P4170), DAPI (D8417), Polybrene (H9268), Puromycin (P8833) were from Sigma (St. Louis, MO, USA), LysoTracker Red (L7528), MitoTracker Green (M7514), Prolong Gold antifade reagent (P36934) were from Thermo Fisher Scientific (Rockford, IL, USA), Ciliobrevin D (250401) from Calbiochem (San Diego, CA, USA) and KOD polymerase (71661) from Novagen (Madison, WI, USA). All chemicals, unless otherwise specified, were obtained from Sigma (St. Louis, MO, USA).

### RNA Interference

HeLa cells were transfected with 1μM each of control siRNA, siRNF167_1, siRNF167_2 separately using RNAimax (13778-075, Thermo Fisher Scientific, Rockford, IL, USA) in reduced serum-containing media (Opti-MEM, 5185-034, GIBCO-Thermo Fisher Scientific, Waltham, MA, USA). The complex was removed after 8-10 hours of transfection and cells were allowed to grow in complete media. siRNA oligos were purchased from Eurogentec (Seraing, Belgium) along with control siRNA (SR-CL000-005). The sequences of siRNAs as follows: RNF167_1, 5’ AAG CAG AGG GAC UGG GUC UUU 3’; RNF167_2, 5’ GGG ACU GGG UCU UCA CUU CUU. Cells were either treated with acetate Ringer’s solution (10 mM Glucose, 70 mM Sodium acetate, 5mM KCl, 80 mM NaCl, 2mM NaH_2_PO_4_, 2mM CaCl_2_, 1mM MgCl_2_, and 10 mM HEPES, final pH 6.9) for two hours (32) or harvested after 24 hours of transfection for immunostaining and qRT-PCR respectively.

### Immunofluorescence and Confocal microscopy

HeLa cells were transfected with RNF167 constructs as mentioned in the figure legends. Briefly, cells were transfected at 18-20 hours of plating (∼60% confluency) using Lipofectamine 3000. Post transfection (24 hours), cells were treated with acetate Ringer’s solution (pH 6.9) for two hours. Cells were fixed in 4% PFA in PHEM buffer (60 mM Pipes, 10 mM EGTA, 25 mM HEPES, and 2 mM MgCl2, final pH 6.8). Immunostaining was performed as previously described (32). Briefly, cells were permeabilized in blocking solution (0.2% saponin + 5% FBS in PHEM buffer) for 30 minutes. Cells were washed in 1X PBS, followed by one-hour incubation with the primary antibody in staining buffer (0.2% saponin in PHEM buffer). These cells were further incubated with a secondary antibody in the staining buffer for 30 minutes. Cells were washed in 1X PBS to remove unbounded antibodies and nuclear staining was performed using 10 μg/ml DAPI for one minute, washed in 1X PBS and mounted in ProLong Gold antifade reagent. All incubations are performed at room temperature unless otherwise mentioned.

To detect the luminal part of LAMP1 in the plasma membrane, cells seeded on glass coverslips were treated with acetate Ringer’s solution (pH 6.9) for two hours. These cells were either treated with DMSO or 5μM ionomycin for 10 minutes and then immediately transferred on to the ice. Immunostaining was performed as previously described (18). Briefly, cells were incubated with anti-LAMP1 (mouse, 1:50) for 30 minutes, 4°C, followed by fixation using 4% PFA in 1X PBS for 10 minutes and incubation with secondary antibody (1:1000) for 1 hour. Next, the cells were stained with 10 μg/ml DAPI and mounted in ProLong Gold antifade reagent.

For live-cell imaging, cells were transfected with the indicated constructs on glass-bottom dishes (MatTek Corporation, P35G-1.0-20-C). Post transfection (24 hrs), imaging dish was loaded into a sealed live-cell imaging chamber (37°C and 5% CO2) for imaging. Single-plane confocal images were acquired using a confocal laser scanning microscope (Leica TCS SP5 II, Wetzlar, Germany) equipped with 63X 1.4 NA oil immersion objective. For quantifications, images were imported into ImageJ (National Institutes of Health, Bethesda, MD, USA) software.

### Quantification of lysosomal distribution and Colocalization analysis

Lysosomal distribution was calculated in terms of the perinuclear index (PNI) (32). Briefly, average LAMP1 intensities were calculated from the whole cell (I _total_), within 5µm distance from the nuclear surface (I _perinuclear_) and more than 10 µm distance from the nuclear surface (I _peripheral_). These intensities were normalized as I_<5_= (I _perinuclear_ / I _total_)-100 and I_>10_= (I _peripheral_ / I _total_)-100 and the PNI was calculated as the difference between I_<5_ and I_>10_ (PNI= I_<5_-I_>10_).

Line Profiling: Line profiling was performed as described earlier (33). The average intensity of LAMP1 positive vesicles was calculated from three independent lines drawn from the nuclear surface. For all three lines, intensities were grouped into three categories, <5µm, 5-15 µm, and >15µm and fractional intensities of each of these regions were calculated using total intensity of that particular line. These fractional values were plotted against the distance from the nuclear surface.

Pearson’s correlation coefficient: Pearson’s Correlation coefficient (PC) was determined using the JACoP plugin of ImageJ (National Institutes of Health, Bethesda, MD, USA) software. PC was calculated using single plane images with threshold adjusted to 17-255 for LAMP1 and 20-255 for RNF167 (default setting: 0-255).

### Plasma membrane repair assays

Plasma membrane repair assays were performed as described earlier (34, 35) and listed below. Immunofluorescence: HeLa cells stably expressing empty vector, RNF167 WT and variants were treated with acetate Ringer’s solution (pH 6.9) for two hours. Cells were washed with ice-cold HBSS thrice and followed by 200 ng/ml Streptolysin-O (SLO) treatment for five minutes on ice. SLO containing buffer was replaced with HBSS or HBSS containing 4 mM CaCl_2_ at 37^ο^C to allow plasma membrane resealing and incubated further for 10 min at 37^ο^C. Next, nuclear staining was performed using 50 μg/ml propidium iodide (PI) for one minute followed by three washes in HBSS at 37ᵒC. Cells were fixed in 4% PFA for 10 minutes. These cells were further washed, incubated with 10 μg/ml DAPI and mounted in ProLong Gold antifade reagent. A minimum of 100 cells from each field was used for calculating the percentage of PI-positive cells in each group.

FACS analysis: HeLa cells stably expressing vector, RNF167 WT and other variants were treated as mentioned above. After resealing, cells were trypsinized immediately. Cell pellets were washed with flow cytometer buffer (1% FBS and 2mM EDTA in PBS) and resuspended it in 250 μl flow cytometer buffer. Before analysis on a BD accuriC6 system (Beckman Coulter, Fullerton, CA, USA), cells were also stained with 50 μg/ml PI. At least 10,000 cells were used for analysis. The percentage of PMR was calculated as described earlier (35).

### RNA isolation and relative quantitative real time-PCR

Total RNA was extracted using the RNeasy Mini Kit (74104, Qiagen, Valencia, CA, USA) following the manufacturer’s recommendation. 1μg of RNA was reverse transcribed using oligo-DT and SuperScript III First-Strand Synthesis System (18080051, Thermo Fisher Scientific, Rockford, IL, USA). Relative qRT-PCR was performed in a total volume of 25 µl with 12.5 µl SYBER GREEN master mix (1725121, BioRad Laboratories, Hercules, CA, USA), 1µl gene-specific primer mix, 5µl cDNA and 6.5 µl sterile water. GAPDH was used to normalize the expression of RNF167 in all the samples used. The normalized values were expressed as relative to control siRNA (considering mRNA level of RNF167 in control siRNA cells as 100%). The primers used for quantitative PCR are listed below:

RNF167-FP-5’-CTGTGGTTGTGGCCGCTGTGCTGTGG-3’

RNF167-RP-5’-CTGCAGGCATTGTCTGGGTGAGCCTCC-3’

GAPDH-FP-5’-ATGGGGAAGGTGAAGGTCGGAGTCAAC-3’

GAPDH-RP-5’-GATCACAAGCTTCCCGTTCTCAGCCTTGAC-3’

### Nested PCR

cDNA prepared from different cell lines (ZR-75, COLO 205, SUM159, HCC1937, MIA PaCa, HeLa, and MCF 10A) were subjected to two rounds of PCR (nested PCR) with the following primers:

FP-5’-GTTTGAAGGTCTCGCGAGATCG-3’ and

RP-5’-CCAGGATTTACGGGTCTTCCGAC-3’

The purified product from the first set of PCR was used as a template for the second step of PCR with the following primers: FP-5’-TCCCACCCAGCTCCACTAAACG-3’and

RP-5’-GCTGGAAGGTCTGCAAAGTCC-3’. Amplicons of expected sizes were purified and confirmed by sequencing.

### Multiple sequence alignment

Amino acid sequences of different isoforms of RNF167 were retrieved from NCBI database with the following IDs (NP_056343.1, NP_001307294.1, NP_001357233.1, NP_001357236.1, NP_001357237.1, NP_001357242.1).These sequences were aligned using Kalign (36) with gap open penalty = 11, gap extension penalty = 0.85 and terminal gap penalty = 0.45. Pictorial representations of the isoforms are based on the multiple sequence alignment.

### Statistical analysis

Data were collected from at least three independent experiments and are shown as mean ± SEM. *P* values were calculated using either two-tailed unpaired Student’s *t* test (for two parameters) or one-way ANOVA or two-way ANOVA (for comparing multiple groups) followed by Tukey’s test as mentioned in the figure legends with significance levels set at α= 0.05. ns-not significant, \**P* < 0.05, \*\**P* < 0.01, \*\*\**P* < 0.001, and \*\*\*\**P* < 0.0001. All statistical analyses were performed in Graph Pad Prism 8 (Version 8.2.0).

## Results

### Identification and Differential subcellular localization of RNF167 isoforms

RNF167, a ubiquitin ligase of 350 amino acids is an endosomal/lysosomal associated protein, with a C-terminal RING domain, N-terminal signal peptide (SP) followed by a protease associated (PA) domain and a transmembrane (TM) domain. However, in the NCBI database, we noted the presence of multiple RNF167 variants of different lengths (Fig.S1.A), suggesting the existence of multiple RNF167 isoforms. Interestingly, a few of the transcripts, particularly, RNF167-b form, lacks the region coding for the first 35 amino acids, raising the possibility that this particular form may differ from the full-length RNF167 (RNF167-a) in its localization to the membrane vesicles and lysosomal associated function (Fig.1A). To test such a possibility first we assessed the expression of RNF167-b. Total RNA from multiple human cell lines were isolated and nested PCR using primers to amplify amplicons of 334-bp and 138-bp for RNF167-a and RNF167-b isoforms was performed (Fig.1 B). Strong band corresponding to 334-bp expected from the expression of RNF167-a was observed in all the cells that were tested. However, some of the cell lines also resulted in an expected size of the 138-bp band, representing the expression of RNF167-b. To confirm that the 138-bp fragment was indeed amplified from RNF167-b, the DNA bands from colon cancer COLO 205 cell lines were extracted, sequenced and confirmed to contain sequences corresponding to RNF167-b (Fig.S1.B). These results established the expression of a RNF167 variant without a predicted signal sequence.

**Figure 1.**
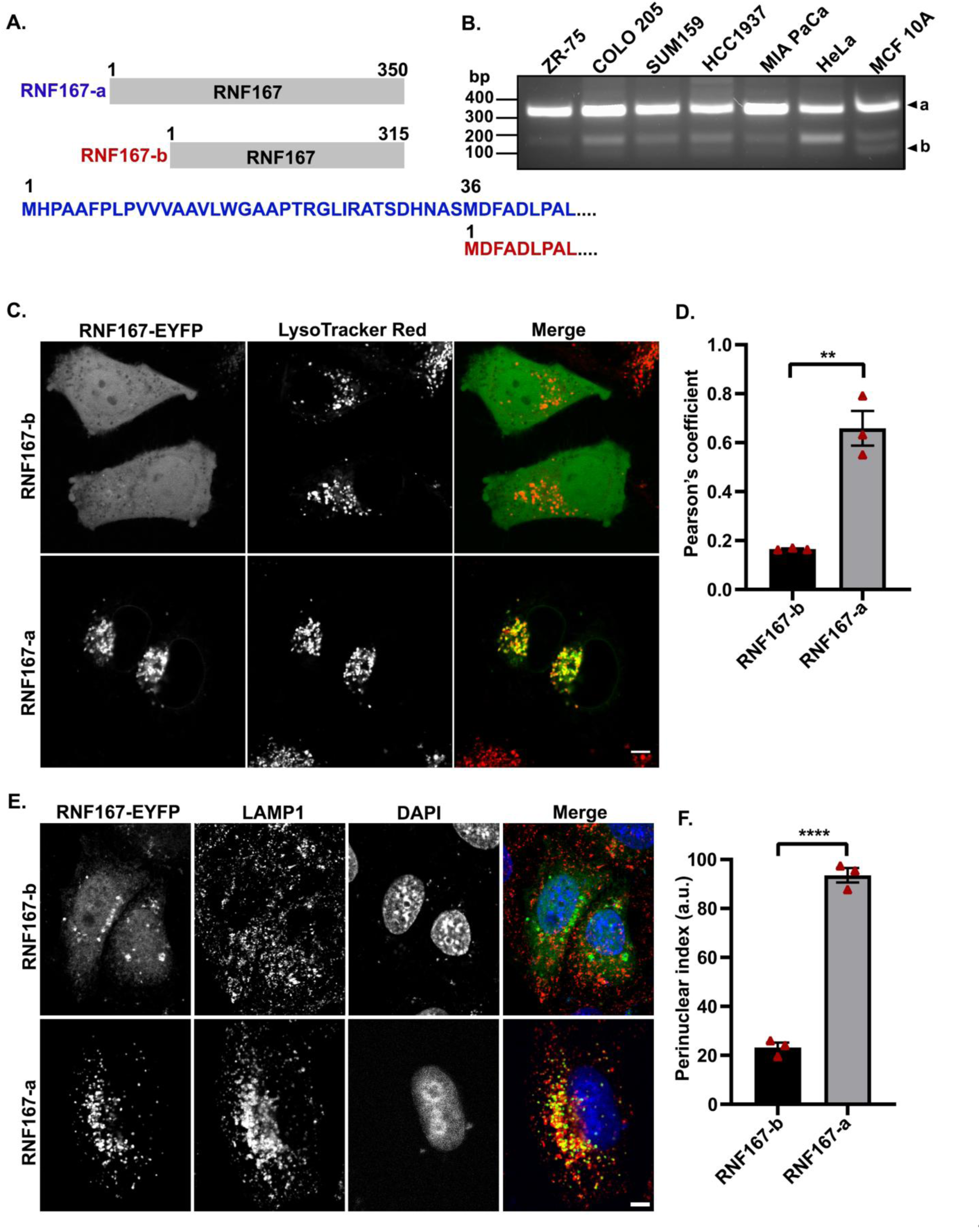
Identification and Differential subcellular localization of RNF167 isoforms. **A)** Schematic representation of two different isoforms of RNF167. **B)** cDNA prepared from indicated cell lines was analyzed by nested PCR and the products were separated on 1.5% agarose gel. The expected amplicon size of RNF167-a (334-bp) and RNF167-b (138-bp) are indicated. **C)** Representative images of HeLa cells transfected with the indicated constructs (green) and stained with 100 mM LysoTracker Red for 30 minutes. **D)** Lysosomal association of isoforms was assessed by measuring the Pearson’s coefficient (n=3, 30 cells per experiment). **E)** Representative images of HeLa cells expressing EYFP tagged RNF167-a and RNF167-b. These cells were treated with acetate Ringer’s solution (pH 6.9) for two hours followed by immunostaining with antibodies against LAMP1 and GFP. **F)** Quantification of lysosomal distribution was done by measuring PNI (n=3, 15-20 cells per experiment). Data represent mean ± SEM. Statistical significance was calculated using Student *t* test (** P<0.01, ****P<0.0001). Scale bars, 5 µm.

To investigate the significance of putative SP, for subcellular localization of RNF167 and its function, we first generated a C-terminal EYFP tagged RNF167-b (RNF167Δ1-35-EYFP) and RNF167-a (full-length), and evaluated their expression pattern in HeLa cells. Interestingly, RNF167-b variant, unlike the RNF167-a which showed punctate intracellular staining, exhibited diffuse cytosolic staining – indicative of requirement of signal peptide for lysosomal localization. Treatment of HeLa cells transfected with RNF167-EYFP isoforms with LysoTracker Red, a lysosomal marker, resulted in a distinct co-localization of RNF167-a and lysosomes, whereas there was a minimal association of RNF167-b with lysosomes (Fig. 1C). The quantification of colocalization and associated Pearson’s coefficient values shown in Fig.1D, revealed the requirement of a signal peptide for RNF167 localization to lysosomes. Interestingly, signal peptide of RNF167, (1–35)-EYFP exhibited diffuse cytosolic staining. This indicates that SP is essential but not sufficient for lysosomal localization of the protein (Fig.S1.C).

As expression of RNF167 is reported to result in perinuclear clustering of lysosomes upon acetate Ringer’s treatment (24), we examined the effect of acetate Ringer’s solution treatment on lysosomal clustering in RNF167-b expressing cells. HeLa cells transfected with plasmids carrying cDNA for RNF167-a or RNF167-b were treated with acetate Ringer’s solution (pH 6.9) and stained with anti-LAMP1 (a lysosomal marker) and anti-GFP antibodies. We noted that consistent with its inability to localize to lysosomal compartments, cells expressing RNF167-b showed substantially more dispersed lysosomes than those cells expressing RNF167-a (Fig. 1E). To independently validate these findings, we assessed the distribution of lysosomes in cells by calculating the perinuclear index (PNI), a widely accepted method to quantitate lysosomal positioning (32). The PNI of RNF167-b was −17.25 as compared to 93.57 of RNF167-a, revealing a regulatory role of RNF167-b variant in lysosomal positioning (Fig.1F).

### RNF167 expression effects positioning of lysosomes but not other endocytic compartments

To determine the specificity of RNF167 mediated lysosomal positioning, we assessed the positioning of various intracellular compartments like endosomes and mitochondria using organelle specific markers; EEA1 (early endosomes), Rab11 (recycling endosomes), mTFP-lysosomes-20 (lysosomes) and MitoTracker Green (mitochondria) in cells expressing RNF167-mRFP and treated with acetate Ringer’s solution. Treatment of the cells with acetate Ringer’s solution is reported to affect mainly lysosomes, not other organelles (6). As expected, we found that mTFP-lysosomes-20 accumulates in the perinuclear region of RNF167 expressing cells, but the distribution of other organelles remains unaffected (Fig. 2A). Collectively these findings suggest that RNF167 status might be a major modifier of lysosomal positioning which is important for various physiological processes, such as repair of the damaged plasma membrane and cellular invasion (1).

**Figure 2.**
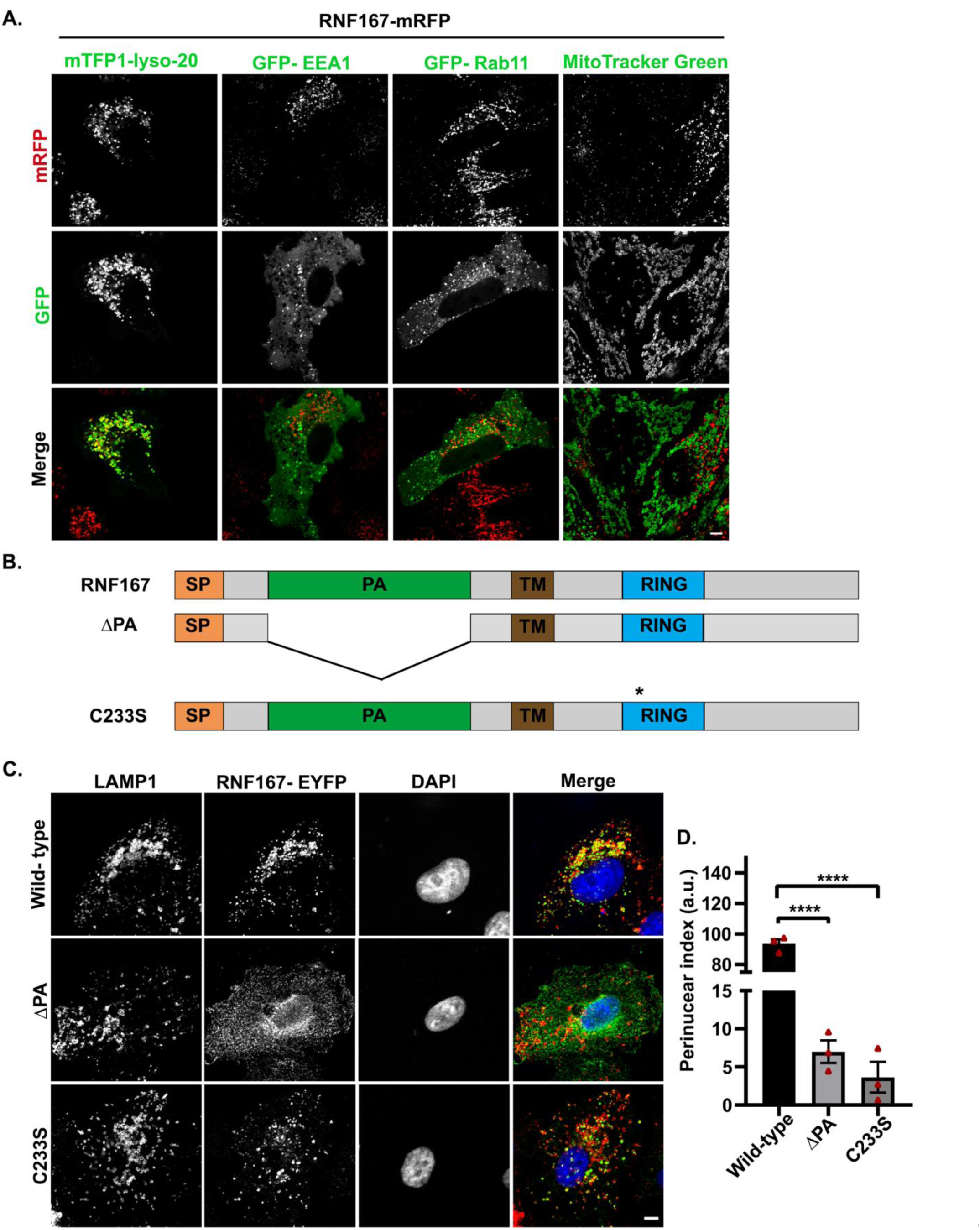
RNF167 mediated clustering is lysosomal specific and is dependent on lysosomal localization and ubiquitin ligase activity of the protein. **A)** HeLa cells stably expressing RNF167-mRFP were either transfected with the indicated constructs to identify different organelle or labeled with MitoTracker Green. Confocal microscopy images of live cells were acquired after two hours of acetate Ringer’s solution treatment (pH 6.9). **B)** Schematic representation of RNF167 variants. **C)** HeLa cells expressing C-terminal EYFP tagged RNF167 wild-type and variants were treated with acetate Ringer’s solution (pH 6.9) for two hours. These cells were stained with antibodies specific for LAMP1 and GFP and images were acquired using confocal microscopy. **D)** Quantification of lysosomal distribution was done by calculating PNI (n=3, 15-20 cells per experiment). Data represent mean ± SEM. Statistical significance was calculated using one-way ANOVA followed by Tukey’s multiple comparison test to compare the means of each sample (****P<0.0001).

### RNF167 mediated lysosomal positioning is dependent on its E3 activity and localization to lysosomes

To establish that RNF167’s localization to the lysosomal compartments is essential for lysosomal clustering, we investigated the structure-function relationship of the PA domain in RNF167. To this end, we have examined the lysosomal positioning in cells expressing RNF167ΔPA (Fig. 2B), a domain reported to be essential for the endosomal localization of the protein. As was the case with RNF167-b (Fig. 1C-E), RNF167ΔPA variant also fails to localize to the lysosomal compartments (Fig. 2C) and induce perinuclear clustering. Moreover, an E3-ligase inactive mutant (RNF167-C233S) with an intact PA domain also failed to cluster lysosomes, suggesting the importance of ubiquitin ligase activity for lysosomal clustering. Quantification of the clustering as measured by the PNI is shown in Fig. 2D.

### Knockdown of RNF167 with siRNA effects lysosomal positioning

To further demonstrate the role of RNF167, we next suppressed the levels of endogenous RNF167 in HeLa cells using selective siRNAs designed to target two independent regions of 3’UTR in RNF167. Transfection of specific siRNAs efficiently downregulated the expression of endogenous RNF167mRNA as compared to the control siRNA (Fig.S1.D). Lysosomal positioning in these cells were examined by immunostaining using anti-LAMP1 antibody and assessed in terms of PNI (Fig. 3A-B). In control siRNA transfected cells, the majority of the lysosomes accumulated in the perinuclear region. Interestingly, depletion of endogenous RNF167 resulted in the dispersal of LAMP1-positive compartments with a significant shift of lysosomes from the perinuclear region to the periphery of the cells (Fig. 3A), indicative of anterograde transport. As expected, the PNI of specific siRNA transfected cells was significantly less as compared to the control siRNA treated cells (Fig. 3B). To further corroborate the results, we assessed lysosomal distribution by line profiling as an established method of choice (33). In control siRNA treated cells, lysosomes were dispersed throughout the cells with a significant population at the perinuclear region (within 5µm distance from the nuclear surface). On the other hand, RNF167 depletion resulted in an increased accumulation of lysosomes at the cell periphery which is defined as more than 15µm from the nuclear surface (Fig. 3C). Collectively, these findings indicate the importance of RNF167 in lysosomal positioning upon extracellular acidification.

**Figure 3.**
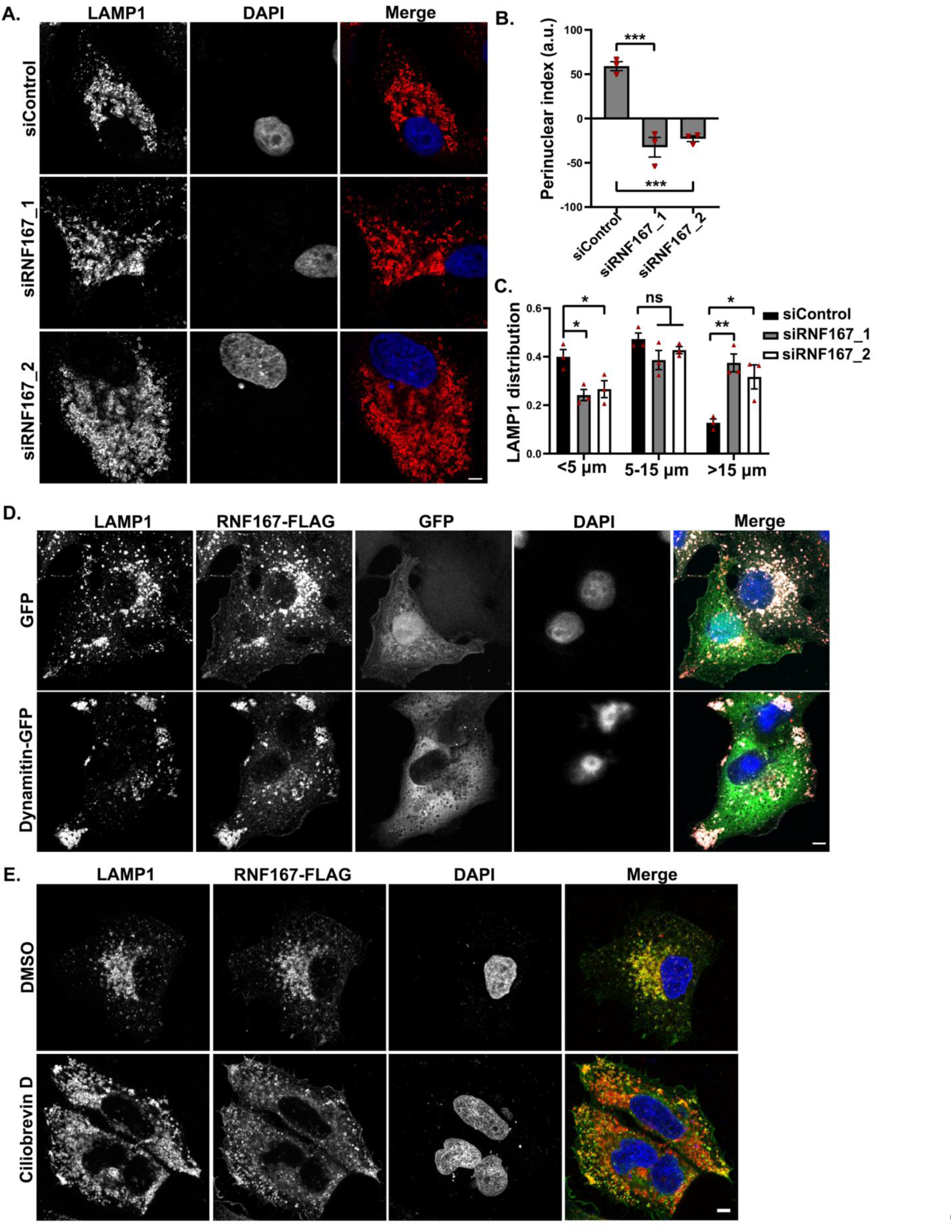
RNF167 promotes dynein mediated retrograde transport. **A)** HeLa cells transiently transfected with control siRNA, or RNF167 specific siRNAs were subjected to two hours of acetate RINGER (pH 6.9) treatment. Lysosomal positioning was analyzed by immunostaining with antibody specific for LAMP1. DAPI represents nuclear staining. **B)** Quantification of lysosomal positioning was done by calculating the perinuclear index. **C)** Lysosomal distribution by line profiling. The fractional intensity of LAMP1 positive compartments was quantified at 0-5 µm, 5-15 µm, and >15 µm. Statistical significance (in **B** and **C**) was calculated using one-way ANOVA followed by Tukey’s multiple comparison test (n=3, 30 cells per experiment, *P<0.05, ** P<0.01, ***P<0.001, ns-not significant). **D)** HeLa cells co-transfected with RNF167-2X FLAG and GFP/Dynamitin-GFP were fixed after acetate Ringer’s solution (pH 6.9) treatment and immunostained with antibodies specific for LAMP1 and FLAG. **E)** HeLa cells transfected with RNF167-2X FLAG and treated with acetate Ringer’s solution (pH 6.9) and 50μM Ciliobrevin D. Immunostaining was performed as in **D**. All data shown are mean ± SEM. Scale bars, 5 µm.

### RNF167 promotes dynein mediated retrograde transport

In mammalian cells, long-range transport of organelles is mediated by microtubule-based motor proteins, including plus end-directed kinesins and minus end-directed dynein complexes (16). Given that RNF167 promotes the accumulation of lysosomes towards the perinuclear region, we hypothesized that RNF167 mediated juxtanuclear positioning of lysosomes might be dynein dependent. To test this hypothesis, we investigated the lysosomal positioning in HeLa cells transfected with RNF167 with or without p50-Dynamitin, overexpression of which is known to disrupt the dynein-dynactin complex (31). As postulated, we found that in RNF167 expressing cells the peripheral pool of LAMP1-positive vesicles is significantly increased in P50-Dynamitin-GFP than GFP alone (Fig. 3D). These observations suggest that RNF167 regulates lysosomal positioning via dynein-dependent retrograde transport mechanism. To further validate the role of dynein motor in this process, experiments were done in the presence of dynein inhibitor, Ciliobrevin D (37, 38). HeLa cells transfected with RNF167 were treated with Ciliobrevin D for two hours before assaying for the lysosomal clustering. We noticed that RNF167 could effectively cluster the lysosomes in the perinuclear region in control cells, whereas treatment with Ciliobrevin D abrogated the lysosomal clustering (Fig.3E) with an increased number of LAMP1-positive compartments in the cell periphery. These results indicate that RNF167-driven perinuclear clustering is specific for lysosomes and dynein-dependent.

### RNF167 regulates lysosomal exocytosis

To understand the physiological significance of RNF167, we focused on the regulation of lysosomal exocytosis in RNF167-expressing cells. Eukaryotic cells reseal the damages on their plasma membrane by a process known as lysosomal exocytosis. Such plasma membrane damages are frequently observed in the cells upon exposure to mechanical stress or during metastasis (18, 39). Damage to the plasma membrane leads to increased intracellular Ca^2+^ ions, lysosomal sensing of elevated intracellular Ca^2+^ and movement of lysosomes towards the plasma membrane in an anterograde fashion to help to repair the damage (40).

We investigated the involvement of RNF167 in lysosomal exocytosis caused by treatment with ionomycin, a known enhancer of intracellular Ca^2+^ level (35, 41) for 10 minutes after 2 hours of acetate Ringer’s solution treatment (pH 6.9). Exocytosis was measured by staining for the LAMP1 on the plasma membrane using luminal epitope-specific anti-LAMP1 antibody. As expected, the cell surface expressions of LAMP1 in control HeLa cells were found to be significantly increased upon ionomycin treatment (1.33% vs. 86%). In contrast, cells stably expressing RNF167 showed a substantial reduction in the surface expression of LAMP1 compared to the control cells under the same condition, 3.33 % as compared to 86% in control cells (Fig. 4A-B). Conversely, cells in which RNF167 was depleted exhibited an increased cell surface expression of LAMP1 (Fig. 4C-D), indicative of exocytosis. Together, these findings suggest that RNF167 influences lysosomal exocytosis by regulating the retrograde movement of lysosomes.

**Figure 4.**
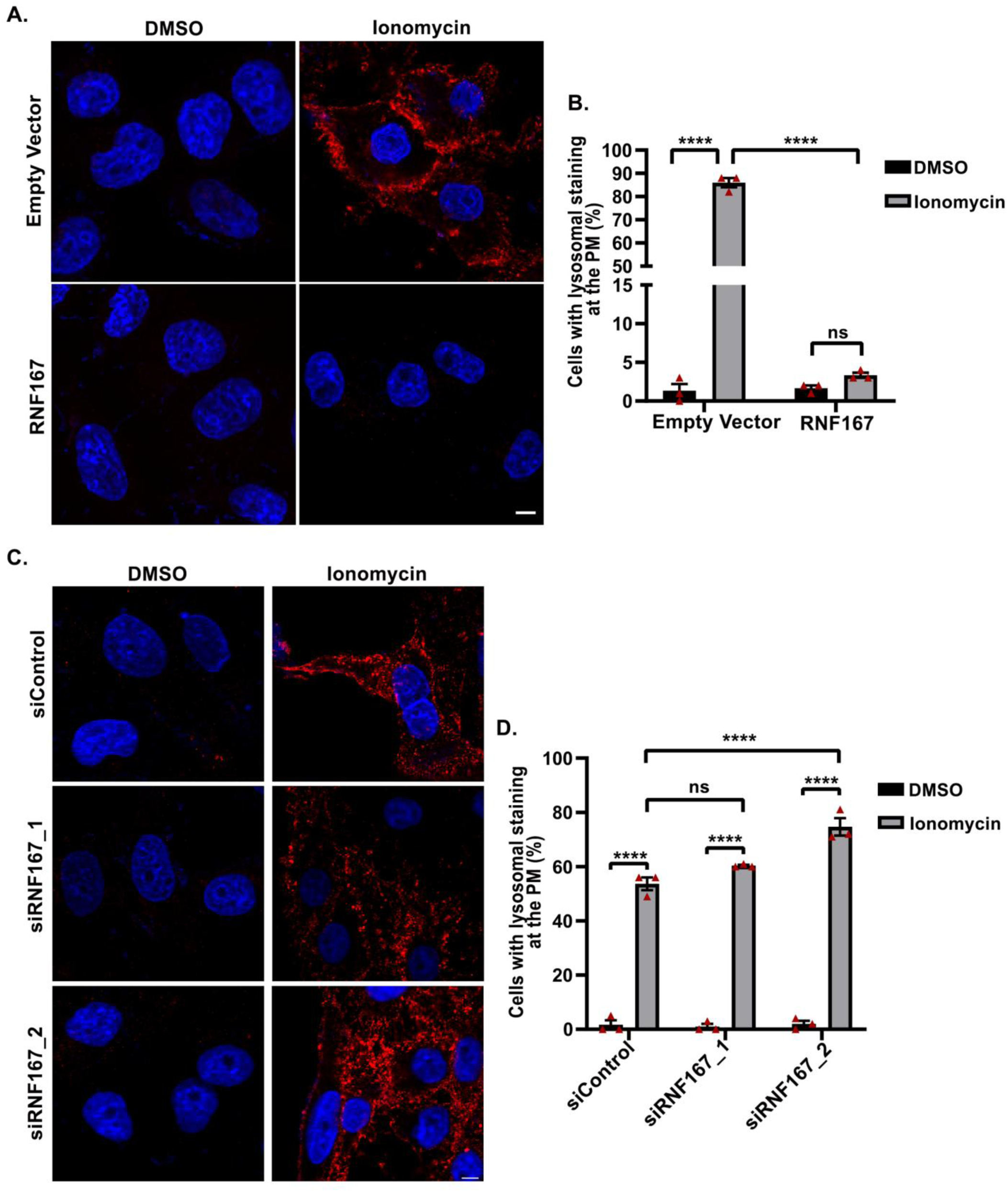
RNF167 regulates lysosomal exocytosis. **A)** HeLa cells stably expressing empty vector or RNF167 were treated with acetate Ringer’s solution (pH 6.9) for two hours. Cells were incubated with 5μM ionomycin for 10 minutes at 37°C and immunostained for luminal epitope of LAMP1 using anti-LAMP1 antibody. **B)** To quantify the surface expression of LAMP1 as a measure of lysosomal exocytosis, a minimum of 65 cells was counted per experiment (n=3). **C)** HeLa cells transfected with control or RNF167specific siRNAs were immunostained for the luminal epitope of LAMP1 as described in **A**. **D)** Quantification of cells for surface expression of LAMP1 described in **C**. A minimum of 55 cells were counted per experiment (n=3). All data shown are mean ± SEM. Statistical significance was verified using two-way ANOVA followed by Tukey’s multiple comparison test (****P<0.0001, ns-not significant).

### Naturally occurring mutations in RNF167 perturb its function

Given the importance of RNF167 for perinuclear clustering of lysosomes and lysosomal exocytosis, we next investigated the effect of a set of naturally occurring mutations identified in RNF167 from diverse cancer-types (http://cancer.sanger.ac.uk). We assessed the ability of tumor associated (TA) mutants of RNF167 (K97N, R200Q, C268R, R277L, and L329P) in regulating lysosomal clustering and exocytosis. Results presented as Fig. 5A indicated that effect of R200Q, C268R, R277L and L329P mutant versions of RNF167 on lysosomal clustering was minimal. However, RNF167-K97N with a missense mutation in the PA domain exhibited a distinct cytosolic expression and also prevented the perinuclear clustering of lysosomes. This suggests that at-least; one of naturally occurring mutation in human tumors affects the lysosomal clustering (Fig. 5B). Since lysosomal distribution is known to affect tumor progression and metastasis, we further investigated the effect of these mutations on lysosomal exocytosis. Consistent with the results shown in Fig. 5 A and B, only K97N but not other mutations or wild-type show a substantial increase in surface expression of LAMP1 (Fig. 5C-D). Because K97N is unable to induce perinuclear clustering, more number of lysosomes are available for docking and fuse with the plasma membrane as evidenced by the increased surface expression of LAMP1.This indicates that the unhindered anterograde movement of lysosomes is an added advantage to the tumor cells, mainly because these lysosomes can promote tumor progression by enhanced lysosomal exocytosis. Given that lysosomal clustering affects plasma membrane resealing as shown with Rab3a depletion (35), we sought to determine the effect of RNF167 and its variants on plasma membrane resealing.

**Figure 5.**
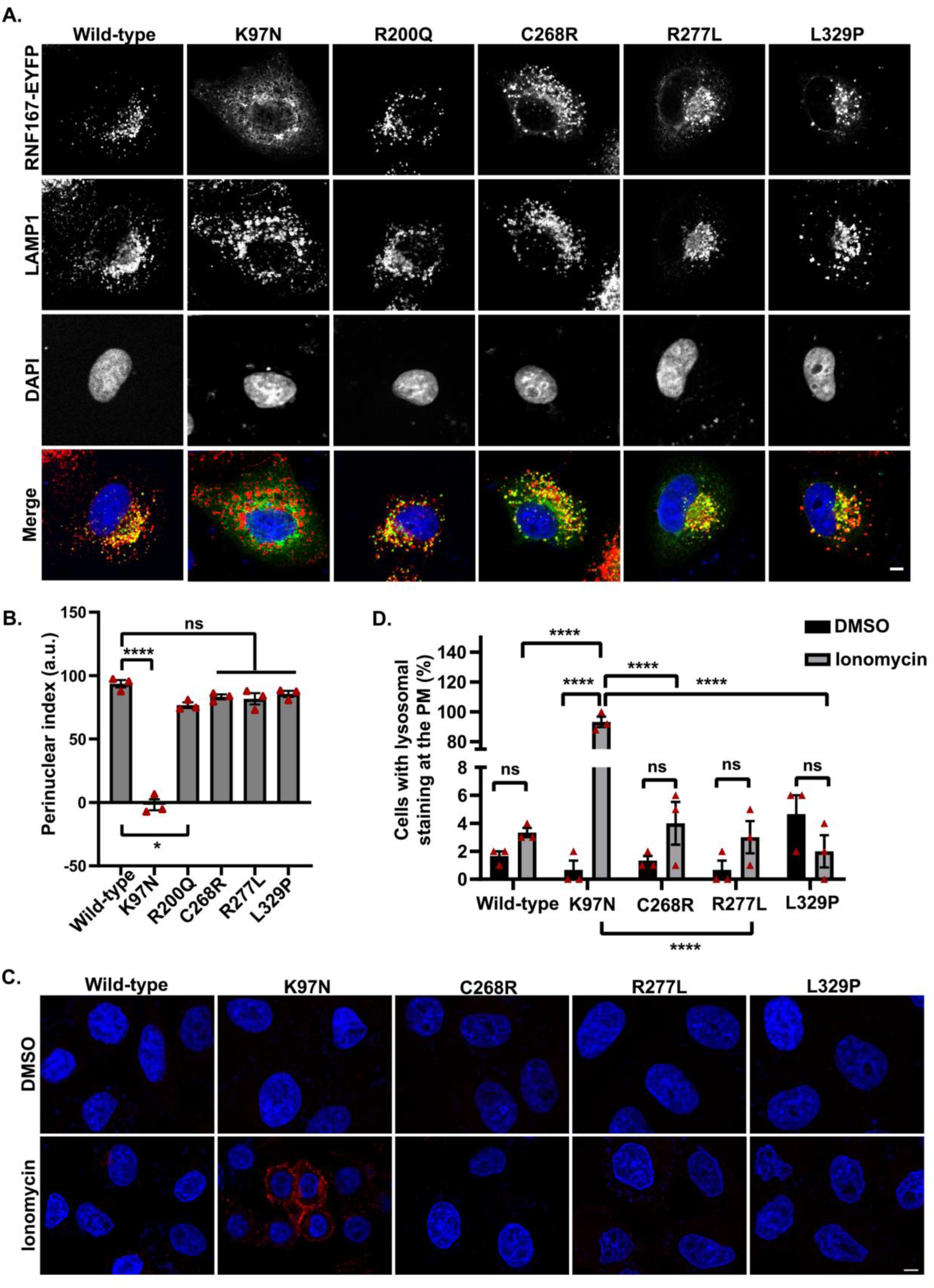
Naturally occurring mutations in RNF167 perturb its function. **A)** HeLa cells expressing RNF167 wild-type or tumor-associated mutants were treated with acetate Ringer’s solution (pH 6.9) for two hours. These cells were fixed and immunostained with antibodies specific for LAMP1 and GFP. **B)** Quantification of lysosomal distribution was done by measuring PNI (n=3, 15-18 cells per experiment). Statistical significance was verified using one-way ANOVA, followed by Tukey’s multiple comparison test (*P<0.05, ****P<0.0001, ns-not significant). **C)** HeLa cells stably expressing RNF167 wild-type or tumor-associated mutants were treated with acetate Ringer’s solution (pH 6.9) for two hours followed by 5μM ionomycin for 10 minutes at 37°C. The presence of LAMP1 on the plasma membrane was detected using luminal epitope-specific anti-LAMP1 antibody. **D)** Quantification of surface expression of LAMP1 described in **C**. A minimum of 180 cells was counted from three independent experiments and the statistical significance was calculated using two-way ANOVA followed by Tukey’s multiple comparison test (****P<0.0001, ns-not significant). All data shown are mean ± SEM. Scale bars, 5μm.

### Cells expressing tumor-associated mutant RNF167-K97N, efficiently repair the plasma membrane

We further examined the effect of RNF167-K69N-mediated lysosomal exocytosis on the ability of cells to reseal their plasma membrane. HeLa cells stably expressing the control vector or RNF167 or RNF167-K97N were treated with acetate Ringer’s solution (pH 6.9) for two hours followed by 200ng/ml Streptolysin-O (SLO) for five minutes. SLO, a bacterial toxin, induces pore formation in the plasma membrane, and is commonly used to study membrane resealing (34). Briefly, cells were allowed to reseal the damage by incubating them in HBSS with or without Ca^2+^ at 37°C for 10 minutes. The efficiency of resealing was measured by propidium iodide (PI) staining as PI is impermeable to cells with the intact plasma membrane. As established earlier (42), the percentage of PI staining in control cells upon Ca^2+^ treatment was much less compared to those without Ca^2+^ confirming the requirement of Ca^2+^ dependent lysosomal exocytosis for maintaining membrane integrity. Consistent with lower LAMP1 surface expression (Fig. 4A-B), cells expressing RNF167 WT cells were unable to repair the membrane resulting in a substantial increase in PI-positive cells, even in the presence of Ca^2+^, and thus, emphasizing the importance of lysosomal exocytosis in maintaining plasma membrane integrity and its regulation by RNF167 (Fig.6A-B). Importantly, cells expressing RNF167-K97N, a localization defective mutant with more dispersed lysosomes was able to reseal the damage more effectively than wild-type expressing cells, leading to lower PI staining (Fig. 6C-D). This enhanced resealing was found only in RNF167-K97N expressing cells but not in cells expressing other tumor-associated RNF167 mutants that were tested. These ineffective mutants showed lysosomal clustering in the perinuclear region (Fig. 5A). This suggests that lysosomal exocytosis might represent an important way to confer a survival advantage to cancer cells, in addition to other processes. Consistent with the above results, increased plasma membrane resealing in cells expressing RNF167-K97N as compared to WT was also confirmed by FACS analysis (Fig. 6E).

**Figure 6.**
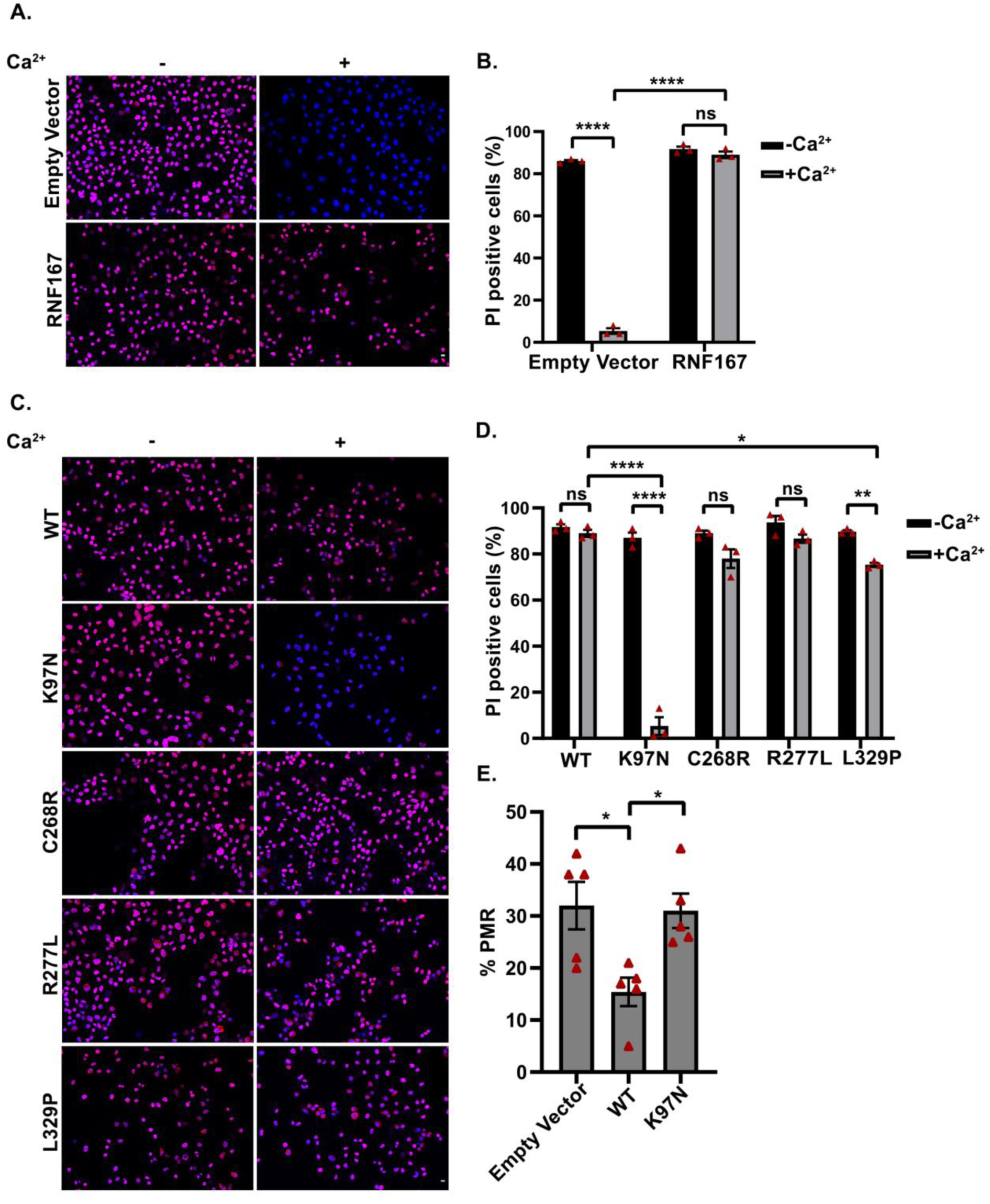
RNF167-K97N, efficiently repair the plasma membrane. **A)** Representative confocal merge images of propidium iodide (PI) stained HeLa cells stably expressing empty vector or RNF167. Cells were treated with acetate Ringer’s solution (pH 6.9) for two hours followed by 200 ng/ml streptolysin-O (SLO) for five minutes on ice and allowed to reseal in the presence or absence of Ca^2+^. **B)** Quantification of percentage PI-positive cells described in **A**. A minimum of 5500 cells was enumerated from three independent experiments and the statistical significance was calculated using two-way ANOVA followed by Tukey’s multiple comparison test (*P<0.05, ****P<0.0001, ns-not significant). **C)** Confocal images of PI stained HeLa cells stably expressing RNF167 wild-type or tumor-associated mutants were treated as mentioned in **A**. **D)** Quantification of percentage PI-positive cells described in **C.** A minimum of 2400 cells was counted from three independent experiments. Statistical significance was calculated using two-way ANOVA followed by Tukey’s multiple comparison test (*P<0.05, ****P<0.0001, ns-not significant). **E)** Percentage of plasma membrane resealing (PMR) in HeLa cells stably expressing empty vector, or RNF167 wild-type, or K97N. Cells were treated as described in **A** and PI-positive cells were measured by flow cytometry (n=5, 10,000 events were analyzed per experiment). Statistical significance was verified by one-way ANOVA followed by Tukey’s multiple comparison test (* P<0.05). All data shown are mean ± SEM. Scale bars, 10 µm.

### Differential regulation of lysosomal exocytosis by isoform b

As shown above (Fig. 1C-D) RNF167-b isoform, a localization defective variant of RNF167 is unable to associate with lysosomes as well as induce perinuclear clustering of lysosomes. To determine whether the naturally occurring shorter version of RNF167 affects the lysosomal exocytosis, we compared the cell surface expression of LAMP1 in HeLa cells stably expressing the RNF167-a or RNF167-b isoforms. Cells were treated with 5µM ionomycin at 37°C for 10 minutes after 2 hours of acetate Ringer’s solution (pH 6.9) treatment. Cells were immediately transferred to ice, stained with the luminal epitope-specific anti-LAMP1 antibody. As expected, ionomycin treatment could not enhance the surface expression of LAMP1 in cells expressing RNF167-a isoform; whereas the plasma membrane staining of LAMP1 in cells expressing RNF167-b isoform exhibited a significant increase under the same experimental conditions.

We noticed about a 16 fold increase in surface expression of LAMP1 in cells stably expressing RNF167-b isoform as compared to cells expressing RNF167-a isoform (Fig.7A-B). Having found an increased surface expression of LAMP1 in RNF167-b isoform expressing cells, we proceeded to measure the plasma membrane resealing capacity by PI staining as described above. As expected, cells expressing RNF167-a isoform had a strong impairment in plasma membrane resealing. In contrast, RNF167-b isoform expressing cells displayed an efficient resealing, as assessed by low PI-positive cells in the presence of extracellular Ca^2+^ (Fig.7C-D). RNF167-b isoform had about nine-fold less PI-positive cells when compared to cells expressing RNF167-a isoform (9.66% PI-positive cells in b and 89% in a). These findings indicate that cells stably expressing RNF167-b isoform has increased plasma membrane resealing by Ca^2+^ dependent lysosomal exocytosis upon extracellular acidification.

**Figure 7.**
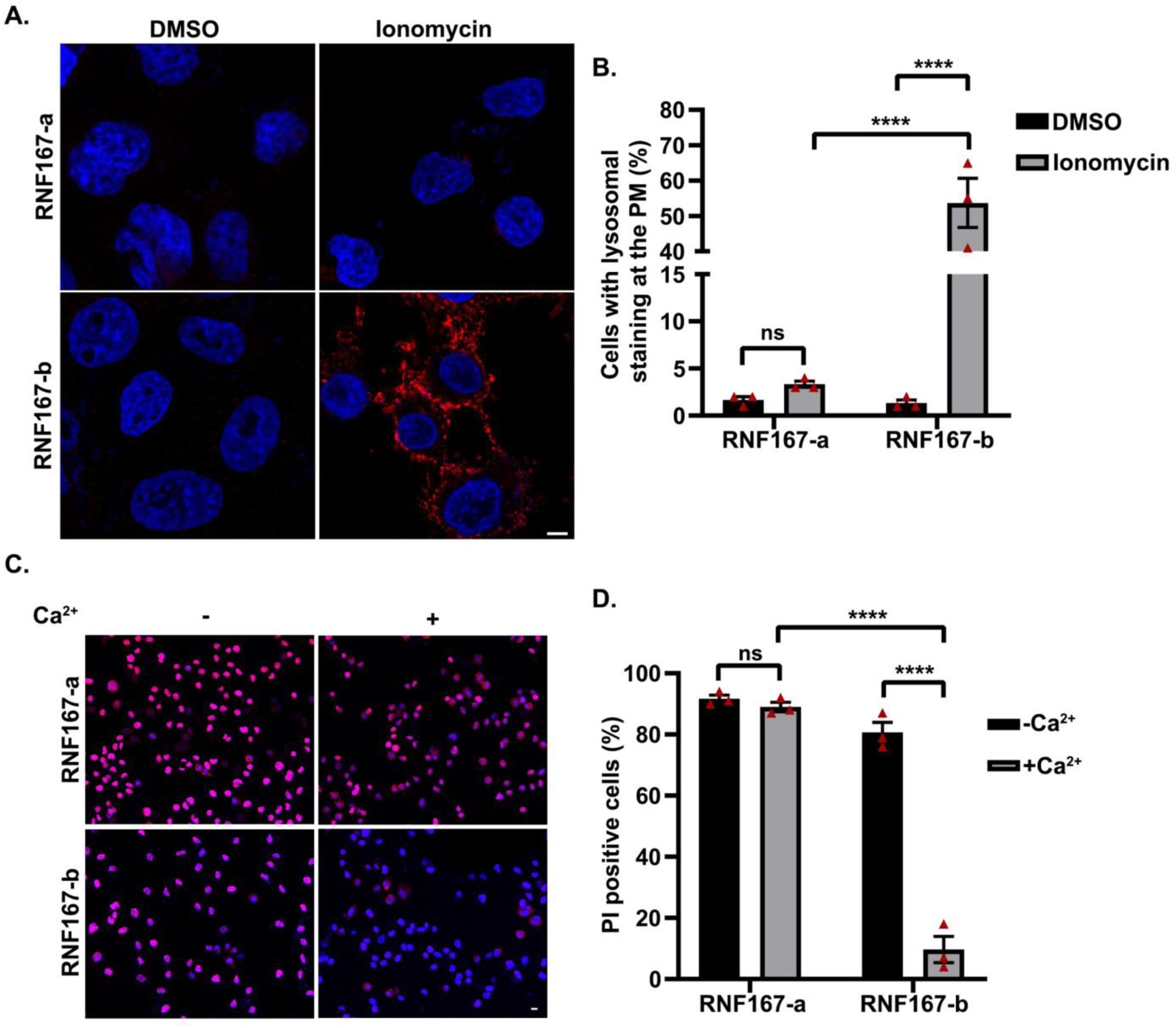
Differential regulation of lysosomal exocytosis by isoform b. **A)** HeLa cells stably expressing either RNF167-a or RNF167-b were treated with acetate Ringer’s solution (pH 6.9) for two hours followed by 5 µM ionomycin for 10 minutes at 37°C. Cells were stained with luminal epitope-specific anti-LAMP1 for 30 minutes on ice. **B)** Quantification of surface expression of LAMP1 described in **A**. A minimum of 70 cells was counted per experiment (n=3). Scale bar-5μm. **C)** HeLa cells stably expressing either RNF167-a or RNF167-b were treated with acetate Ringer’s solution (pH 6.9) for two hours followed by 200 ng/ml SLO for 5 minutes on ice. These cells were allowed to reseal the damage in the presence or absence of Ca^2+^. **D)** Quantification of PI-positive cells from merged images of HeLa cells stained with DAPI and PI after the SLO treatment. A minimum of 3200 cells was counted from three independent experiments. All data shown are mean ± SEM and the statistical significance was verified using two-way ANOVA followed by Tukey’s multiple comparison test (****P<0.0001, ns-not significant). Scale bar, 10 µm.

We also assessed the lysosomal exocytosis in HeLa cells transiently transfected with EYFP tagged RNF167-a and RNF167-b. As expected, cells expressing RNF167-b-EYFP exhibited increased surface expression of LAMP1 upon ionomycin treatment (Fig. S2.B). This is in line with our finding that RNF167-b-EYFP expressing cells have more dispersed lysosomes upon extracellular acidification. Collectively, our results suggest that the two isoforms of RNF167 could affect tumor progression differently by promoting opposing functions in the regulation of lysosomal positioning, exocytosis, and PMR.

## Discussion

Lysosomes undergo long-range intracellular transport along microtubules with the help of motor proteins kinesins and dynein and short-range transport via actin-based myosin motors (2). In this study, we show that RNF167, a lysosomal associated ubiquitin ligase, regulates the intracellular movement of lysosomes. We found that overexpression of RNF167 promotes dynein mediated retrograde transport of lysosomes whereas depletion of endogenous RNF167 results in their anterograde movement. Our findings are further supported by the recent observation that Arl8B, one of the key regulators of lysosomal anterograde movement, is a substrate for RNF167 (24). We identified that the PA domain of RNF167 is required for its localization to the lysosomal compartments and is essential for the perinuclear clustering of lysosomes. We further observed that RNF167 localizes to Rab7 positive endosomes in addition to lysosomes (data not shown) raising the possibility that RNF167 can also affect conventional Rab7-RILP mediated retrograde movement of lysosomes along the microtubules. It would be interesting to elucidate the role of RNF167 in recruiting dynein motors to lysosomes.

Interestingly, in several patient tumor samples, different mutations in RNF167 have been identified. Analysis of the COSMIC database (http://cancer.sanger.ac.uk/cosmic) shows that 67.16% changes are due to missense substitution followed by 19.4% synonymous substitution, 10.45% frameshift deletion and 1.49% each of inframe deletion and nonsense substitutions. RNF167 is a ubiquitin ligase, and mutations in the RING domain can affect its ligase activity. Moreover, mutations in other domains like PA or SP can affect not only the stability of the protein but also localization, and function of the protein. RNF167-V98G, a TA mutant identified from patients with lung cancer, harbors a missense mutation in the PA domain is shown to abrogate localization of the protein to endosomal compartments (22). Because of the cross-talk between lysosomal positioning and plasma membrane resealing and its importance in tumor progression, the effect of RNF167 and its TA mutations on lysosomal exocytosis and PMR processes have been investigated in this paper. Out of the five mutations (K97N, R200Q, C268R, R277L, and L329P) that were tested, one mutation in the PA domain (K97N) was unable to localize to lysosomes and to induce perinuclear clustering. This is consistent with the observations shown in Fig. 2C with RNF167ΔPA, indicating the importance of lysosomal localization for RNF167 function. To our surprise, RNF167-C268R, which harbor a missense mutation in the RING domain still able to induce perinuclear clustering, unlike RNF167-C233S. RNF167-C233S, a ligase inactive mutant that has been reported to block perinuclear clustering of lysosomes (24). To understand these differences, we assessed the lysosomal association of C268R and C233S, two distinct ligase inactive mutants of RNF167 by measuring the Pearson’s coefficient. It was indeed evident from the Pearson’s coefficient that C268R had a better lysosomal association as compared to C233S (Fig. S2.A). The differences in clustering may be attributed to their intracellular localization or reasons yet to be found. In conclusion, ubiquitin ligase activity and the lysosomal localization of the protein are essential for regulating lysosomal positioning. Perinuclear clustering of lysosomes is not affected by mutants R200Q, R277L and L329P.

Intracellular mobility of lysosomes plays an important role in tumor cell invasion and metastasis (43). Cells are prone to plasma membrane damage during the invasion of the extracellular matrix which tumor cells partly offset by enhanced lysosomal exocytosis and that helps in plasma membrane repair (18). Acidic extracellular pH (pHe), a specific feature of the tumor microenvironment, has been shown to stimulate the anterograde movement of lysosomes. Hence it is conceivable that mutations in regulatory molecules of exocytosis favoring anterograde movement would be beneficial to tumor progression. In fact, it is known that tumor cells change the expression level of some of the regulators of anterograde movement for their advantage. For example, skeletal muscle sarcoma cells enhance lysosomal exocytosis by downregulating the expression of NEU1, a sialidase and a known negative regulator of lysosomal exocytosis. Deficiency of NEU1 leads to the accumulation of over sialylated LAMP1 and these actively participate in lysosomal exocytosis (20).

To our knowledge, no TA mutations in regulators of lysosomal exocytosis have been characterized. In this report, we have investigated the effect of several TA mutations of RNF167 on lysosomal exocytosis and PMR and demonstrated that one of the mutations, K97N, affected lysosomal exocytosis (measured by immunostaining using luminal epitope-specific anti-LAMP1 antibody) and PMR. K97N mutant was reported from kidney tumor samples and the lysine residue at the 97^th^ position is highly conserved. The reason for increased lysosomal exocytosis and PMR exhibited by K97N expressing cells is because of the dispersed lysosomes compared to the WT expressing cells. The effect on lysosomal exocytosis is unique for K97N but not for other mutations (C268R, R277L, and L32P) that have been tested. Also, K97N, not other mutants showed defects in lysosomal localization. It is possible that these mutations of RNF167 can influence yet to be discovered pathways without affecting lysosomal localization resulting in tumor progression. It would be interesting to look at lysosomal positioning and exocytosis in primary cells derived from tumor samples of patients with mutations in RNF167.

Interestingly, our findings also reveal an important role for RNF167 isoform. We found that isoform b results in splicing of an exon that code for 1-35 amino acids, signal peptide. We demonstrated that signal peptide is also required for lysosomal localization in addition to the PA domain as RNF167-b showed cytosolic localization. We have confirmed the expression of RNF167-b in COLO-205 cells by sequencing the amplified PCR products. Moreover, we found that RNF167-b, unlike RNF167-a, enhances lysosomal exocytosis and PMR. We provide evidence for functional differences between the two variants. Differential expression of isoforms with antagonistic functions has been associated with tumor progression. For example, Intersectin1 (ITSN1) is a highly conserved scaffold protein and exists in two isoforms referred to as long isoform (ITSN1-L) and short isoform (ITSN1-S). ITSN1-S promotes glioma development whereas ITSN1-L inhibits glioma progression both in vivo and in vitro. This opposing function of ITSN1-S and ITSN1-L is highly regulated by alternative splicing (44). Similarly, alternative splicing of CD99 leads to two isoforms, full-length (CD99wt) and a truncated short form (CD99sh) (45, 46). CD99 is a transmembrane glycoprotein that plays a crucial role in cell adhesion, apoptosis, differentiation of T cells and thymocytes (45, 47), etc. Studies from osteosarcoma and prostate cancer cells revealed that the full-length form inhibits anchorage-dependent growth, migration, and metastasis whereas the shorter form promotes tumor progression (48). In addition to this, alternative splicing of Bcl-x produces two isoforms, Bcl-x (L) and Bcl-x (S). Bcl-x (L) promoted cell growth and proliferation and inhibited programmed cell death, while Bcl-x (S) promoted cell apoptosis (49, 50).

Hence it is conceivable that isoforms of RNF167 could also play a major role in maintaining homeostasis at the organism level. In summary, our study identifies RNF167 as an important regulator in Ca^2+^ dependent lysosomal exocytosis and plasma membrane resealing. We demonstrated that K97N, one of the tumor-associated mutants of RNF167 is unable to localize to the lysosomal compartments and cells expressing this mutant efficiently repair the plasma membrane by enhanced lysosomal exocytosis. We also show that a localization defective variant of RNF167, isoform b, lacking 1-35 amino acids, also failed to cluster lysosomes in the perinuclear region and shows similar phenotype as K97N. Our data reveal new clues for the regulation of lysosomal exocytosis and tumor progression.

## Abbreviations

PMR: Plasma membrane resealing
PI: Propidium iodide
SLO: Streptolysin-O
DAPI: 4’, 6-Diamidine-2’-phenylindole dihydrochloride
PFA: Paraformaldehyde
PNI: Perinuclear index
WT: Wild-type
TA: Tumor-associated mutants

## Acknowledgements

The authors great fully acknowledge the following people for their generous gift of reagents: Dr. Roberto Botelho, Ryerson University (Dynamitin-GFP); Dr. Ruth H Palmer, University of Gothenburg (C-terminal GFP tagged RNF167-C268R, RNF167-R277L, RNF167-L329P, and RNF167ΔPA). The authors also acknowledge technical help from Keerthi. S. Nair and Fabi Rasheed. SVN and RPN acknowledge financial support from DST-INSPIRE for Ph.D. fellowship and undergraduate fellowship respectively. NDN acknowledge financial support from IISER Thiruvananthapuram (IISER TVM). This work was partly supported by the Department of Science and Technology-Science and Engineering Research Board (DST-SERB) grant (EMR/2016/008048) and IISER TVM awarded to SMS.

## Author Contributions

SVN, NDN and SMS conceived and designed the study. SVN, NDN and RPN performed the experiments and collected the data. RK contributed RNA from different cell lines. SVN, NDN and SMS analyzed the results. SVN, RK and SMS prepared the manuscript.

## Supplemental material

**Figure S1.**
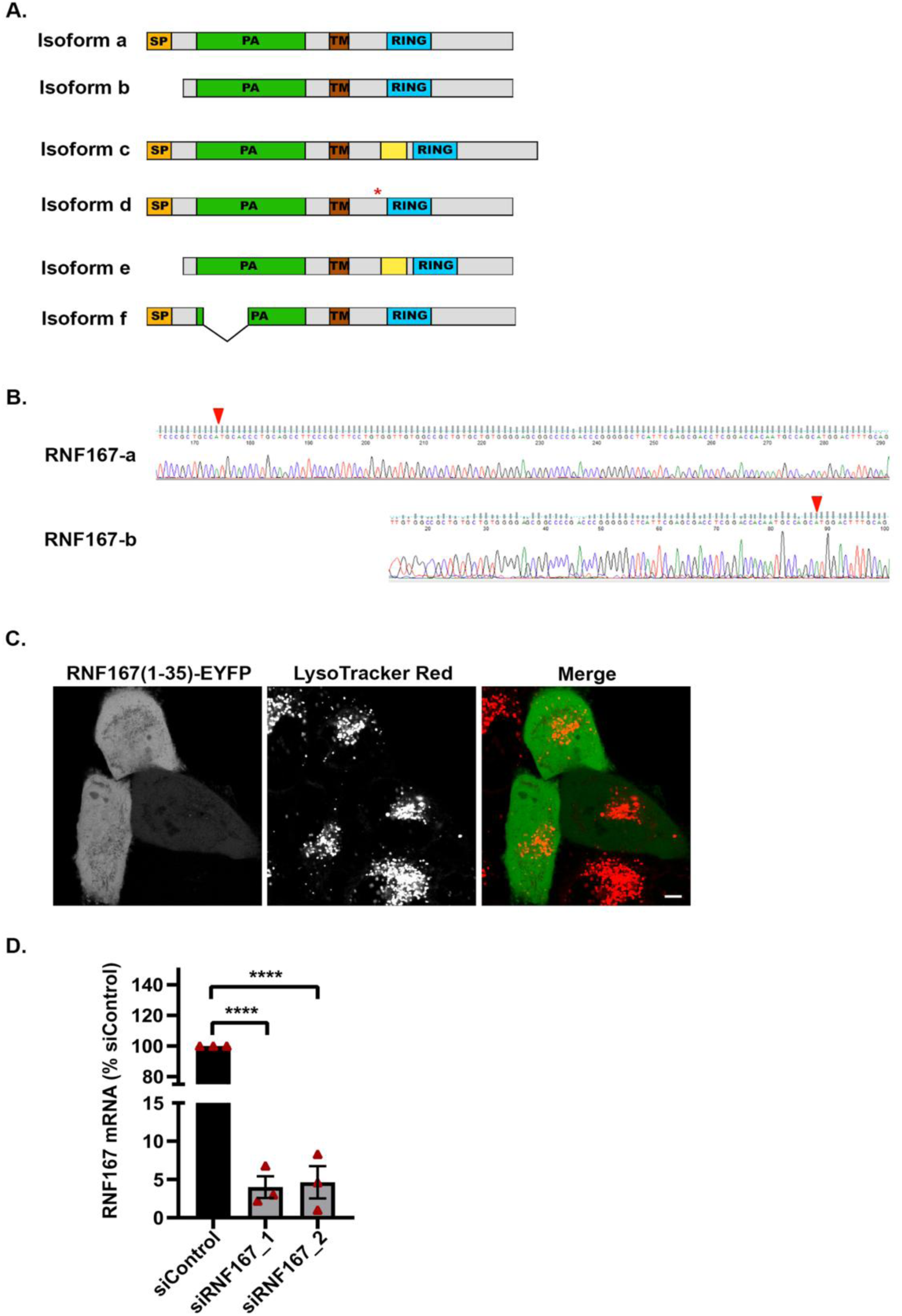
Information supporting Figs. 1 and 3. **A)** Pictorial representation of different isoforms of RNF167. SP-Signal peptide, PA-Protease associated domain, TM-Transmembrane domain, and RING domain. In comparison to isoform a, other isoforms have either an additional 24 amino acids length uncharacterized region or deletion of 42 amino acids within the PA domain, or deletion of SP or an amino acid at 224^th^ position. **B)** Sequencing result of RNF167-a and RNF167-b specific PCR products obtained from COLO 205 cell line. Starting codon in both forms is highlighted in red. **C)** RNF167 (1-35)-EYFP is cytosolic and is unable to associate with LysoTracker positive compartments. HeLa cells transfected with RNF167 (1-35)-EYFP were labeled with 100 mM LysoTracker Red for 30 minutes and subjected to live cell imaging. Scale bar, 5μm. **D)** Effectiveness of siRNA specific for RNF167 was analyzed by qRT-PCR. HeLa cells were transfected with 1µM each of control siRNA, or siRNF167_1 or siRNF167_2 separately, and mRNA levels of RNF167 was measured. The data was normalized with GAPDH and represented as relative to control siRNA (n=3). Data represent mean ± SEM. Statistical significance was calculated using one-way ANOVA followed by Tukey’s multiple comparison test (****P<0.0001).

**Figure S2.**
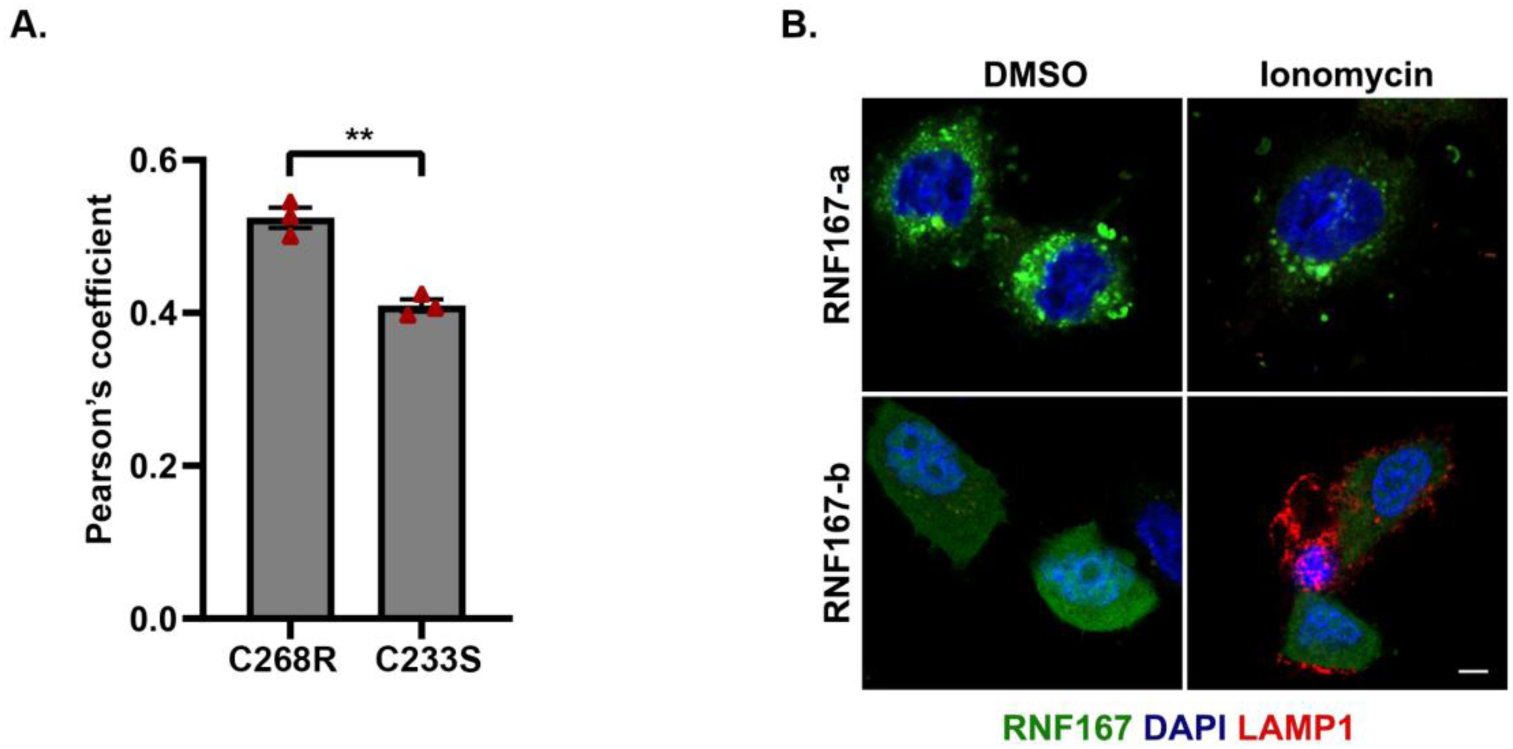
Information supporting Figs. 5 and 7. **A)** Association of RNF167-C233S and C268R variants with LAMP1 positive vesicles. Colocalization of LAMP1 with two distinct ligase inactive mutants of RNF167 (C233S and C268R) was assessed by measuring the Pearson’s coefficient (n=3, 20 cells per experiment). Data represent mean ± SEM. Statistical significance was calculated using Student *t* test (** P<0.01). **B)** Enhanced lysosomal exocytosis by RNF167-b. HeLa cells transfected with EYFP tagged RNF167-a and RNF167-b separately. 24 hours of post-transfection, cells were treated with acetate Ringer’s solution (pH 6.9) for two hours followed by 5μM ionomycin for 10 minutes at 37°C. The presence of LAMP1 on the plasma membrane was detected using luminal epitope-specific anti-LAMP1 antibody. Scale bar, 5μm.

